# Plasma exosomal miRNA analysis of Alzheimer’s disease reveals the dysfunction of a neural network

**DOI:** 10.1101/2021.04.12.439575

**Authors:** Yuzhe Sun, Zhen Hefu, Wang Lifang, Benchao Li, Song Zhijie, Yan Deng, Liu Zhili, Jiahong Ding, Tao Li, Wenwei Zhang, Nie Chao, Shuang Rong

## Abstract

Exosomal microRNA (miRNA) is an emerging source for biomarkers of Alzheimer’s disease (AD). Here, we profiled miRNA expression in AD, mild cognitive impairment (MCI), and controls. The assessment and validation of differentially expressed miRNA represented their potential to be novel biomarkers for AD and MCI. We conducted 13 co-expression networks and a miRNA network module linked to neural function emerged as the most significantly associated with AD diagnosis. The conservation analysis revealed the M1 was highly preserved in controls but dysfunction in AD and MCI. The module pattern between MCI and NC was similar, but significantly differed from AD, suggesting that the neural network regulated by miRNA changed during the mild cognitive stage, and the total miRNA expression altered in AD stage. Additionally, 24 out of 26 M1 hub-miRNAs were derived from brain tissue, and 15 had been reported as AD biomarkers. We consequently proposed the other 11 miRNAs could play important roles in AD. Our study highlights that co-expression network analysis can provide a new path for finding novel biomarkers.

## Introduction

Alzheimer’s disease (AD), caused by neurodegeneration, is second-most common in elderly populations ^1^. The slowly progressing neuropathological changes, such as amyloid β (Aβ) deposition, initiate at the early stage of dementia, which is called mild cognitive impairment (MCI) ^2^. It is the pathological clinical status between cognitively normal and AD ^3^. Lengthening the early stage of dementia can delay the onset of AD pathology, highlighting the need to identify biomarkers for the pathological and molecular changes in the early stage of dementia ^4^. The early diagnosis for AD is challenging because neurodegenerative phenotypes are usually accompanied by MCI, which is difficult to distinguish from normal dementia caused by aging ^5, 6^. Although efforts have been made to identify early-stage biomarkers, the most significant clinical parameters for AD are still Aβ and Tau ^7, 8^.

Extracellular vesicles (EVs) can pass through the blood-brain barrier, making it a valuable source of molecular diagnostics for neurological diseases. In the central neural system, 70% of miRNAs are released from human brain cells and may regulate the transcription of more than one-third of all genes ^9, 10^. Tissue-specific miRNAs can be isolated from enriched extracellular vesicles such as exosomes, the most intensely studied subtype of EVs. In physiological and neurodegenerative brain disorders, enriched miRNA and other cellular components are released from neuronal cells, and spread from cerebrospinal fluid (CSF) to peripheral blood carried by exosomes ^11^. Therefore, exosomes isolated from peripheral blood could represent a promising source of biomarkers for neurodegenerative disease (ND), further improving the detection and prediction accuracy at the pre-clinical stage.

To date, several potential miRNA biomarkers for ND have been examined in blood, plasma, and serum by qPCR or small RNA sequencing, including miR-149-5p, miR-124-3p, miR-9-3p/5p, and miR-125b-5p ^12–15^. A single miRNA usually has several target genes that belong to different functions, while a single gene can be targeted by multiple miRNAs. The complexity of miRNA mediated regulatory mechanism leads to an absence of information on the gene interaction network. Weighted correlation network analysis (WGCNA) is used to find highly correlated expression patterns, and calculate the relationship between selected modules and external sample traits ^16^. The classification of ND is to enhance precision and prediction of early-onset through molecular characterization. Gene expression-based network analysis has been shown to be useful for the prognosis of many neurodegenerative disease subtypes ^17, 18^. Groups of aberrantly expressed genes may work in a modular fashion to promote neurodegeneration, resulting in progression of disease symptoms ^17^. Thus, the deactivation and activation of coexpression networks could also be associated with known biological functions which might uncover new mechanisms underlying the progression of neurodegenerative diseases.

In this study, we compared the exosome-derived miRNA profiles from plasma of normal control (NC), MCI and AD participants. We assessed the diagnostic power of differentially expressed miRNAs for MCI and AD. Co-expression network analysis revealed a network highly correlated with the AD diagnosis. This network was conserved in NC but dysfunction in MCI and AD, very different from the miRNA total patterns, in which NC and MCI were like each other. Tissue-specific analysis showed most of the hub miRNAs in this network were specifically expressed in brain, indicating these miRNAs may originate from the brain and reflect the post-transcriptional regulation in the nervous system.

## Material and Methods

### Participant enrollment

In this study, 158 subjects participated in this study including: 48 AD patients, 48 MCI patients, and 62 NC. Subjects were from communities of Wuhan City, Huangshi City, and Jingmen City in Hubei province, China. All cases were frequency matched by age and sex for utilizing more samples and available information. The diagnosis of AD was based on the criteria of the National Institute on Aging and Alzheimer’s Association (NIA-AA). The diagnosis of MCI was based on a previous study ^19^. The battery of cognitive tests including Mini-Mental State Examination (MMSE), Montreal Cognitive Assessment (MoCA), Clinical Dementia Rating (CDR), Hachinski Ischemic Score (HIS), and Geriatric Depression Scale (GDS) was applied to AD patients. All participants were approved by the BGI and the Medical Ethics Committee of Medical College, Wuhan University of Science and Technology. The standardized questionnaire was employed to obtain the information on demographic characteristics, history of diseases, and lifestyle factors. Written consent was also obtained from all subjects.

### Biochemical measurements

The levels of silent information regulator-1 (SIRT1) protein, interleukin-6 (IL-6), total tau (T-tau), and phosphorylated tau (P-tau), and amyloid β 1-42 (Aβ1-42) in plasma were measured using enzyme-linked immunosorbent assay (ELISA), the coefficients of variation both between and within the plates of the ELISA Kits (Elabscience® Biotechnology Inc., United States) were < 10%.

### Exosomal RNA extraction and small RNA (sRNA) sequencing

Approximate 0.5 mL plasma samples were collected from AD/MCI participants and controls, and stored at −80℃ prior to RNA extraction. Exosomes were isolated from all 500ul plasma using ExoQuick™ plasma prep and exosome precipitation kit (Cat No. EXOQ5TM-1-SBI, Palo Alto, USA), followed by small RNA extraction procedures using miRNeasy Serum/Plasma Kit (Cat No. 217084, QIAGEN, Hilden, Germany). All exosomal isolation and sRNA extraction experiments were carried out strictly according to the manufacturer’s instructions. All sRNA samples were subsequently run on a Bioanalyzer 2100 system (Agilent, Santa Clara, US) using Agilent small RNA kit. RNA integrity number (RIN) was also measured by Bioanalyzer 2100 system. The concentration of sRNA was measured by Nano Drop 2000 (Thermo Scientific). Small RNA sequencing libraries were prepared from total exosomal sRNA with the MGIEasy Small RNA Library Prep Kit (MGI, Shenzhen, China). Pooled libraries were loaded and sequenced on BGISEQ-500 (BGI), and more than 2 Gb of reads were obtained from each library.

### Small RNA data analysis

We acquired an average of 2.2 Gb raw data per sample after small RNA seq. Raw reads were processed by trimming adaptor sequences, and low-quality reads were removed. Clean reads with the size 70-500 Mb per sample were mapped against the reference genome (hg19) using Bowtie 2 ^20^ with 1 mismatch. The matched reads were aligned to mature miRNAs in miRbase version 20^21^, miRNA counts were calculated with perfect-matched reads. The remaining reads were mapped to Rfam database ^22^ to identify and remove protein-coding genes, transfer RNA (tRNA), and ribosomal RNA (rRNA) to predict the novel miRNAs. We used miRDeep2 to predict the novel miRNA from the retained reads^23^. Differential expression was performed using two software EdgeR and DESeq2 ^24, 25^. First, differentially expressed miRNAs were identified separately by EdgeR and DESeq2. Then, the intersection of the two lists was defined as the final differential expressed miRNAs. The expression levels of miRNA were displayed by TPM (transcript per million). The thresholds of significant miRNA expression changes were defined as over ± 1-fold (log_2_) and FDR ≤ 0.05.

R package MultimiR was used to predict targets of miRNA ^26^. Databases including Targetscan ^27^, miRTarBase ^28^, miRDB ^29^, and DIANA-TarBase ^30^ were used in the prediction and an average of ten targets per each miRNA was predicted. Gene ontology (GO) analysis was performed from potential target genes using clusterProfiler ^31^.

### Statistics

We selected 13 differentially expressed miRNAs to assess the diagnostic power for AD and 16 differentially expressed miRNAs to assess the diagnostic power for MCI. Predictive modeling was performed using the R package Caret to combine the predictive power of multiple miRNAs. In detail, logistic regression with internal 10-fold cross-validation was used to develop the model. The area under the receiver operator characteristic curves (AUC) was calculated to assess the power of the model by R package pROC.

### Weighted gene correlation network analysis (WGCNA)

The co-expression network analysis was performed using the R package WGCNA^16^. To construct a weighted coexpression network, expression levels of 158 sRNA-seq samples were used, including AD, MCI, and NC groups. The WGCNA network function was used with the following settings: soft threshold power β= 5, deepSplit = 4, merge cut height of 0.07, minimum module size of 12. Pearson correlations between each miRNA and each module eigengene were performed. After the initial network construction, 13 modules consisting of 12-202 miRNAs were detected. The first principal component (eigengene value) was calculated and considered as representative of each module. We discarded the grey module, which cannot be merged into any other modules, and then correlated the eigengene values of modules with phenotype traits shared by AD, MCI, and NC samples. To characterize the co-expression network, we selected the miRNAs with the highest eigengene values, predicted the targets, and performed functional enrichment analysis using Metascape^32^. Based on the GO annotation of miRNA targets, we then summarized the potential function of each network.

We separately assessed the network preservation in AD, MCI, and NC groups. Using the networks obtained above as the template, Z_summary_ composite preservation scores were calculated to each target group, with 500 permutations. The modulePreservation() function in the WGCNA package was used.

## Results

### Overview of study design and demography of AD, MCI, and control samples

To investigate the miRNA expression profile in AD patients, we collected 158 plasma samples from 48 sporadic AD cases, 48 sporadic MCI cases and 62 NC cases. The detailed patients’ demography is shown in Table 1. There were no differences in the ages or ratios of males/females among the AD, MCI, and NC. The MMSE scores were significantly lower in AD and MCI patients than in NC patients, while the MoCA scores were significantly lower in MCI patients than in NC patients (Table 1, Supplementary Table 1). MoCA and MMSE are important cognitive assessments for AD, significant differences among AD, MCI and NC confirm the rationality of grouping.

Additionally, we collected the existing disease and lifestyle factors of each patient. Education and marriage were also different among three groups which was consistent with the previous studies ^33^. Among existing diseases, hypertension and stroke could raise the risk of developing AD (Table 1). Lifestyle (low education, low body mass index (BMI), marriage status, smoking, less tea-drinking, less entertainment, low reading frequency, less exercise, and less communication with neighbors and children) increases the AD risk (Table 1, Supplementary Table 1).

### Exosomal miRNA expression profiles in AD and MCI patients

To obtain the exosomal miRNA transcriptome, we constructed and sequenced miRNA libraries from plasma exosome samples of AD, MCI, and NC patients. After removal of low-quality reads, we identified 1718 miRNAs. Using DESeq2 and EdgeR software calculation results, 71, 43, 92 differentially expressed miRNAs were identified between AD and NC, MCI and NC, MCI and AD, respectively (FDR > 0.05, *p*-value < 0.01) (Supplementary Table 2)^24, 25^. After removing miRNAs of | log_2_ (fold change) | < 1, 16 and 13 miRNAs were differentially expressed in MCI and AD patients, respectively (Table 2). All but two miRNAs (miR-6891-5p and miR-7975) were down-regulated in AD and MCI patients, indicating an overall decline of exo-miRNA expression in participants with cognitive decline (Table 2, Fig. 1A). Eight miRNAs, including miR-124-3p, miR-9-5p, miR-153-3p, miR-9-3p, miR-338-3p, miR-127-3p, miR-125b-5p, and miR-3529-3p, were down-regulated in both AD and MCI patients, suggesting similar miRNA expression changes in AD and MCI samples (Fig. 1B, Table 2). The down-regulation of miR-9-3p and miR-9-5p is consistent with previous research utilizing autopsied brain tissue^34^. In AD-MCI, 13 miRNAs were increased in AD samples, while only 2 miRNAs were decreased, suggesting that expression levels of some miRNA didn’t alter according to the disease severity (Table 2). In summary, most exo-miRNA exhibited decreased expression levels in both AD-NC and MCI-NC comparisons, and a few miRNAs in AD were increased compared to MCI.

**Figure 1.**
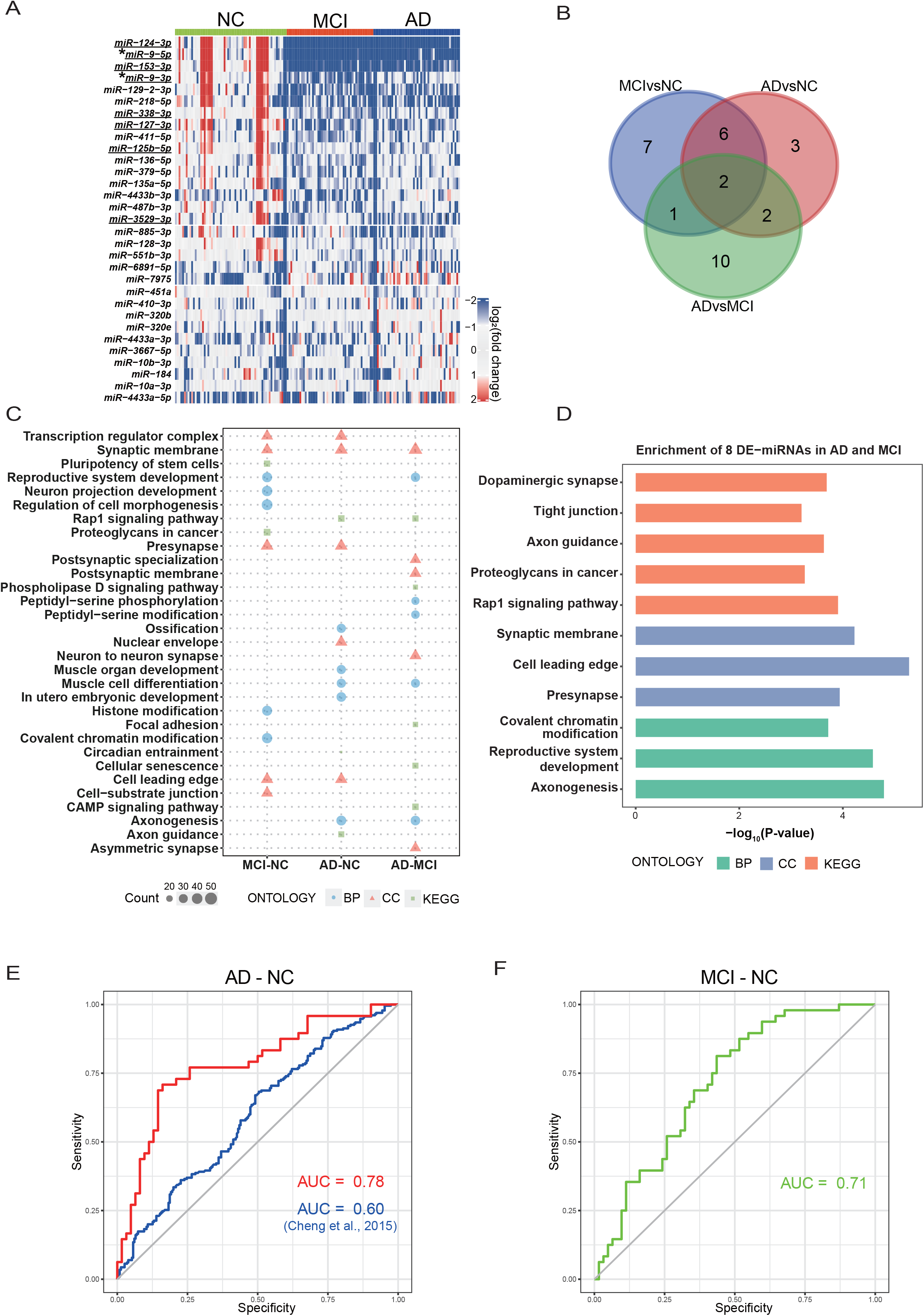
Exosomal miRNA expression profiles, pathway enrichment and miRNA diagnostic prediction in AD and MCI patients. (A) Differentially expressed miRNAs in AD and MCI patients. The expression levels (log_2_) of differential miRNA are shown by two-color heatmap. Underline represents 8 commonly declined miRNAs in AD and MCI. Asterisk (*) represents two differentially expressed miRNAs in three comparisons (AD-NC, MCI-NC, and AD-MCI). (B) Venn diagram of differentially expressed miRNAs in AD-NC, MCI-NC, and AD-MCI comparisons. (C) GO and KEGG enrichment of altered miRNAs. BP: biological process, CC: cellular component. (D) GO and KEGG enrichment of eight declined miRNAs in MCI and AD. (E) ROC analysis shows the area under the curve (AUC) of AD-NC classifier is 0.78 in this study, and 0.60 in a published study^38^. (F) ROC analysis shows the AUC of MCI-NC classifier is 0.71.

Next, we performed target prediction and gene enrichment of target genes using multiMiR and clusterProfiler^26, 31^. Significantly altered AD miRNAs enriched in cellular components of the synapse-related neuron to neuron communication and biological process of axonogenesis by gene ontology (GO) analysis (Fig. 1C), which is consistent with earlier findings that dysfunctional synapses are widely observed in neurons of AD patients^35^. Kyoto Encyclopedia of Genes and Genomes (KEGG) pathway also showed an enrichment in axon guidance in AD patients (Fig. 1C). In MCI group, cellular component such as presynapse, synaptic membrane, and biological process in neuron projection development were found. The same enrichment in synapse-related component and axonogenesis process were observed in the differentially expressed miRNAs between MCI and AD (Fig. 1C). Eight suppressed miRNAs in AD and MCI were enriched in neuronal cells processes, and were predicted to play roles in axon and synapse (Fig. 1D). The above results are supported by previous studies^36, 37^ and imply that the altered expression levels of exo-miRNAs may derive from changes in neuronal exosomes, making them a potential peripheral blood biomarker for early diagnosis.

Furthermore, we examined the predictive power of the differentially expressed miRNAs of MCI-NC and AD-NC comparisons. DESeq2 was used to standardize the miRNA expression levels, and the differentially expressed miRNAs were considered the covariates. The clinical phenotypes were predicted by the logistic regression. Finally, the AUC in the AD-NC comparison is 0.78 (*p* < 0.001), and the AUC in the MCI-NC comparison is 0.71 (*p* < 0.001) (Fig. 1E, Fig. 1F). These results indicate that differential expression of miRNAs in blood neuronal-derived exosomes could be potential diagnostic markers for AD. Additionally, we downloaded exo-miRNA profiles from a previous study^38^. Using the same miRNAs and criteria, AUC in the AD-NC is 0.60, which is much lower than in our prediction (Fig. 1E). It is possible because the smaller sample size (23 NC and 23 AD) and the miRNA extraction (from serum) is different in that research.

### Construction of a consensus AD miRNA coexpression network

To further understand the biological function of plasma exo-miRNA, we generated a coexpression network from the top 1000 abundant miRNAs using a WGCNA algorithm. The networks consisted of 13 co-expressed miRNA modules with similar expression patterns across 158 cases (Fig. 2A). The module sizes ranged from 202 (M12, turquoise) to 12 (M4, salmon). GO analysis of the miRNA module members revealed potential function of 13 modules, encompassing a diverse mix of biological processes and pathways (Fig. 2A, Supplementary Table 3). To assess whether a given coexpression module was related to AD and other patient phenotypes, we correlated module eigengene values of the module miRNA expression level to gender, educational history, cognitive function assessed by MMSE and parameters in a personal health survey, including marriage, medical history, smoking, alcohol and tea drinking, entertainment, reading behavior, exercise frequency, communication with neighbors and children (Fig. 2A). Three modules are significantly correlated with AD diagnosis and cognitive measures: M1 (neural function), M3 (transcription repressor), and M7 (GTPase binding). The M1 neural function module exibits the strongest AD trait correlations (AD diagnosis, *p* = 9 × 10^−5^; MMSE, *p* = 0.004), indicating the miRNAs in M1 module are strongly correlated with AD neuropathology. Vertically, M3 and M7 are correlated with several parameters in the survey, while education level and reading behavior are correlated with more than half of the modules (Fig. 2A). These findings suggest that some biological function of M1, M3 and M7 may be altered during disease progression. It is worth noting that four modules (M2 SMAD binding, M5 calcium-transporting, M6 transcription coregulator, M12 proximal promoter) are correlated with MMSE (*p* < 0.05), indicating that the functions in these modules may also effectively affect cognition.

**Figure 2.**
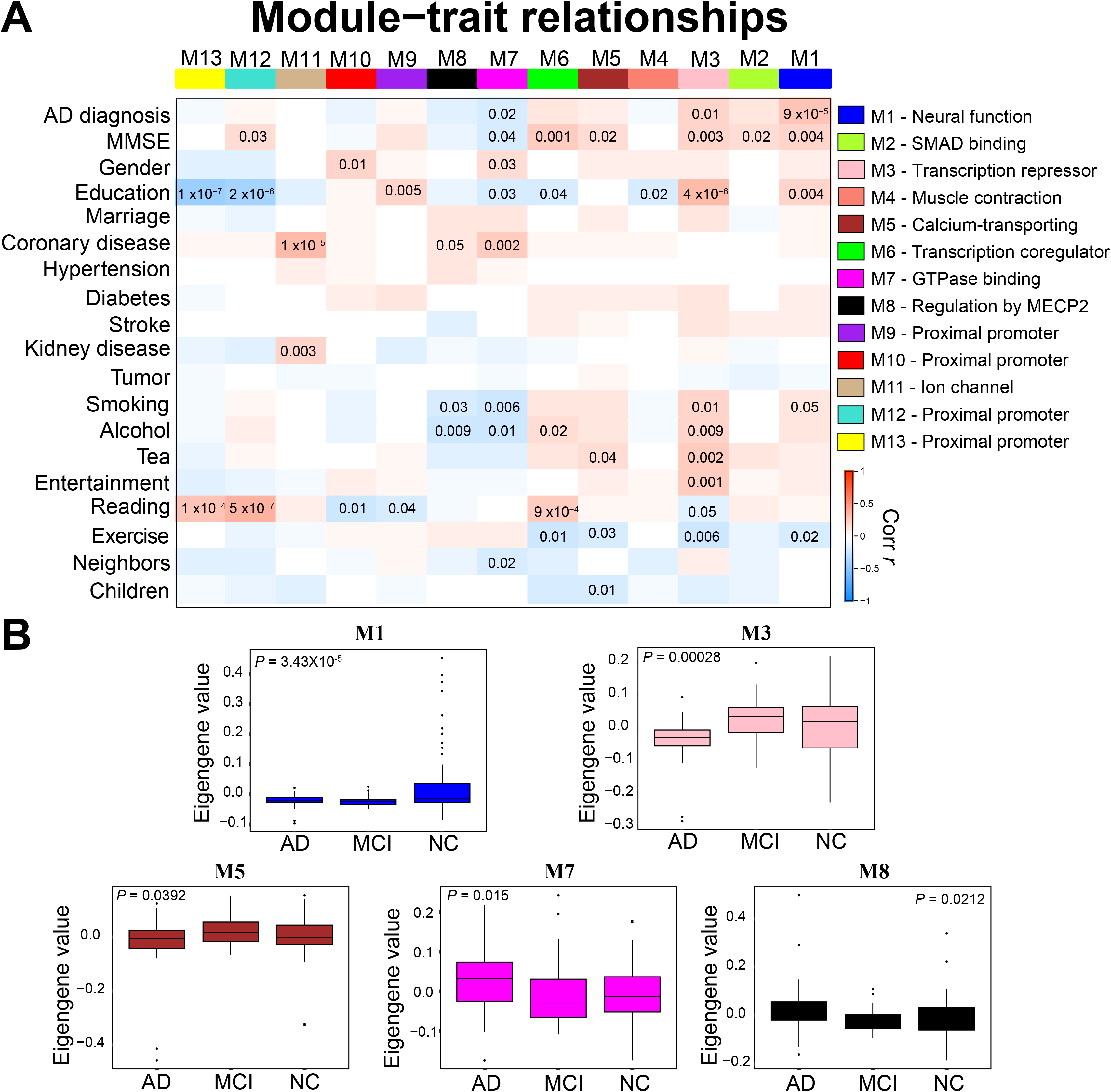
Co-expression network analysis of miRNA profiles in AD, MCI, and NC. (A) Correlation network consisting of 13 modules was generated from top 1000 abundant miRNAs. Module eigengenes were correlated with AD diagnosis, cognitive function (MMSE) and personal surveys. Strength of positive (red) or negative (blue) correlation is shown by two-color heatmap, with *p* < 0.05 values provided. GO analysis of the miRNA targets within each module annotated the biological processes associated with the module. (B) Module eigengene values by case status for 13 modules. Case status is from 158 participants. Differences in eigengene values were calculated by Kruskal–Wallis one-way analysis of variance (ANOVA).

In order to investigate the relationship between the diagnostic classification and the network modules, we summarized the eigengene values (the first principal component of the expression matrix) in each module by case status. Five modules exhibited significant differences among three diagnostic categories, including M1 neural function, M3 transcription repressor, M5 calcium-transporting, M7 GTPase binding, and M8 regulation by MECP2 modules (Fig. 2B). Compared to NC, AD and MCI in M1 were significantly decreased and exhibited the strongest *p* value (*p* = 3.43 × 10^−5^), indicating that M1 neural function module had dramatically changed in MCI stage, before AD (Fig. 2B). In other modules, eigengene values of M3 and M5 decreased in AD and increased in MCI, while eigengene values of M7 and M8 increased in AD and decreased in MCI (Fig. 2B). In general, compared to NC, modules that were increased or decreased in AD were altered in the opposite orientation in MCI, except M1. It suggests that M1 is the most important module representing the AD pathology, and miRNAs in M1 could be potential biomarkers for staging AD progression.

The coexpression networks M1, M3, M5, M7, and M8 contained 105, 25, 86, 20, and 25 miRNAs, respectively (Supplementary Table 4). We calculated the membership of each miRNA (kME) and defined the miRNAs with top 25% kME values in each co-expression module as hub miRNAs. Beta diversity (sample dissimilarity) based on weighted distances was performed using hub miRNAs by case status (Fig. 3A). Unlike the eigengene values of the co-expression modules, diversity of M3 and M5 hub miRNAs revealed significant differences compared to NC, indicating the function of M3 transcription repressor and M5 calcium-transporting changed in the MCI stage (Fig. 3A). Similar patterns of M1 and M8 were found between eigengene values and hub miRNA diversity (Fig. 2B and 3A). The beta diversity patterns of M1, M3, M5 and M8 revealed that the major differences occurred in the MCI stage, suggesting that the post-transcriptional regulation from co-expressed miRNA underwent from disease onset of the progressive phase.

**Figure 3.**
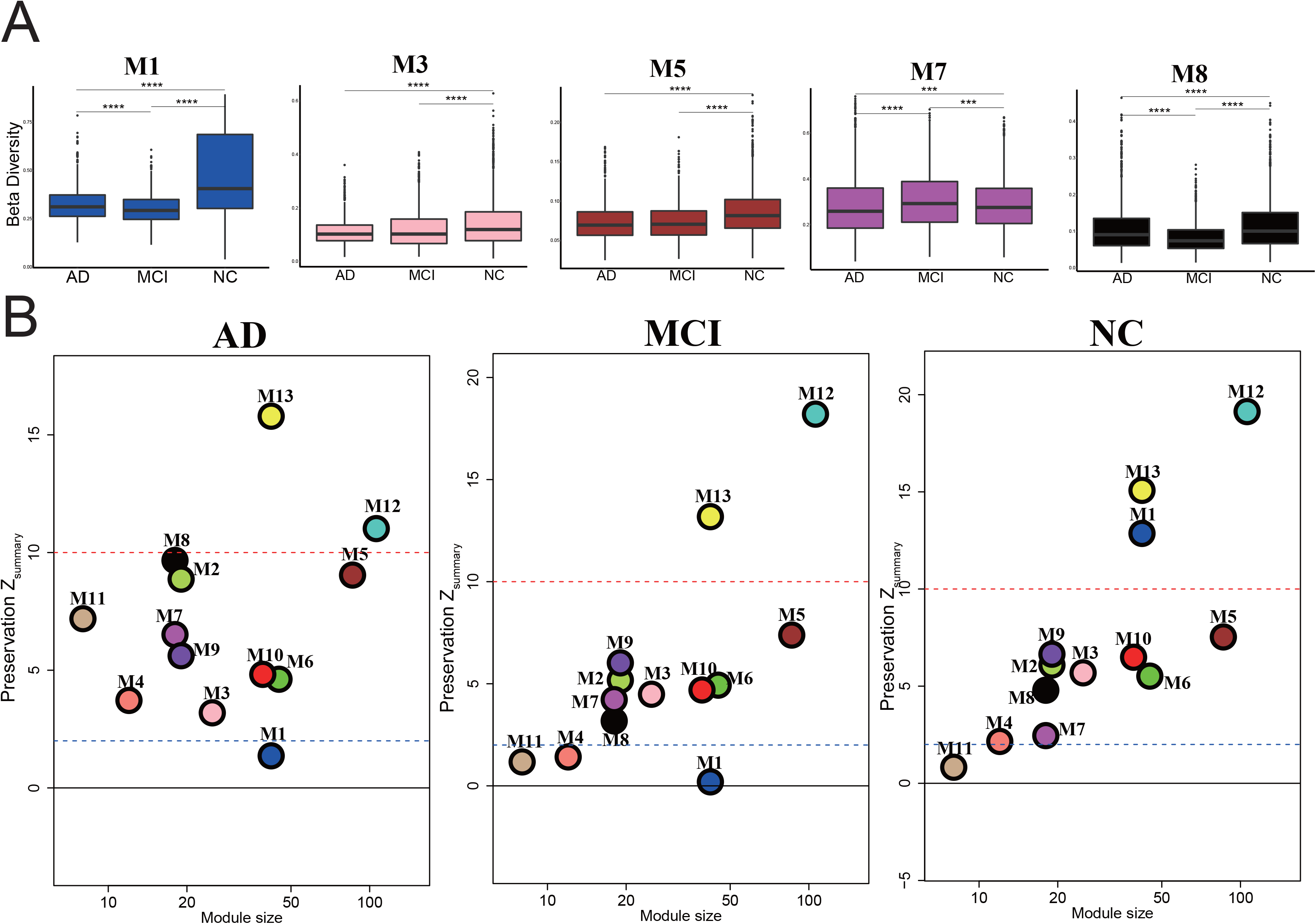
Beta diversity of hub miRNAs and module preservation of AD network. (A) Box plots of intra-group beta diversity based on hub miRNA expression profiles in AD, MCI, and NC (\**p* < 0.05, \*\**p* < 0.01; Wilcoxon rank-sum test). (B) Co-expression network preservation. The dashed blue line indicates a Z_summary_ score of 2 or false discovery rate (FDR) q value < 0.05, above which module preservation was considered statistically significant. The dashed red line indicates a Z_summary_ score of 10 or FDR q value ~1 × 10^−23^.

### The M1 network has the potential to be biomarker for AD early diagnose

To assess the conservation of the network modules in the three groups, we analyzed network module preservation statistics by calculating a composite Z_summary_ score^39^. Derived from 158 samples, patterns of module preservation were similar between MCI and NC, and were different compare to AD (Fig. 3B). It indicates that the miRNA expression will be totally altered after the clinical pathology of AD. Between MCI and NC, there was a major difference: M1 neural function (Fig. 3B). Interestingly, M1 was highly preserved in NC samples (Z_summary_ = 12.9), but the preservation scores were below the threshold in AD and MCI samples (Z_summary_ = 1.3 in AD, Z_summary_ = 0.2 in MCI, the threshold = 2) (Fig. 3B). It indicates that the connectivity inside the M1 network was weakened during the progression of the disease, and this occurred in the MCI stage. Within the 26 hub miRNAs in the M1 network module, 15 were associated with AD or neural function. Meanwhile, 8 differentially expressed miRNAs in AD and MCI were clustered into M1, and 6 of them were hub miRNAs (Table 3). It implies the strong relationship between the M1 network and AD (Table 3).

Besides, M11 ion channel was not preserved in MCI and NC, and M4 muscle contraction was not preserved in MCI (Fig. 3B). M12 and M13 were highly preserved (Z_summary_ > 10) in all three groups, indicating exo-miRNA related proximal promoter could be a house-keeping function that didn’t change greatly in disease progression.

### The connection of AD-specific measurements and the co-expression network

We collected CDR, HIS, GDS, APOE genotype, Systolic/Diastolic pressure, heart rate, blood glucose, Cholesterol, Triglycerides, HDL, and LDL of AD patients. Using the same modules constructed in 158 samples, we assessed the relationship between AD patients’ miRNA expression and phenotypes (Fig. 4A, Supplementary Table 5). Three modules (M1, M2 and M9) had close relationships with both HIS and GDS, while M5 and M7 were closely related to APOE genotype (Fig. 4A). Interestingly, M11 ion channel correlated with several health indicators, such as systolic pressure, heart rate, cholesterol, and HDL (Fig. 4A). It suggests that the regulatory function of M11 exo-miRNA in blood could be a composite index of vascular health. Furthermore, we measured the concentration of SIRT1, IL-6, T-TAU, P-TAU, and Aβ42 proteins of 26 AD patients and observed a significant relationship between M8 and IL-6, *p* = 0.029 (Fig. 4B). M1 revealed a high correlation with SIRT1 but did not show significance, probably because of the small sample size (Fig. 4B).

## Discussion

In this study, we demonstrated that 16 and 13 neural-derived exo-miRNAs were significantly altered in MCI and AD patients, respectively (Fig. 1A and 1B). The diagnostic power of these miRNAs revealed some significance in AD-NC and MCI-NC comparisons (Fig. 1E and 1F). Then, a weighted gene co-expression network analysis was performed and M1 neural function revealed a very high correlation with AD diagnosis (*p* = 9×10^−5^) (Fig. 2A). A module preservation analysis of 13 identified modules showed an overall similarity between the patterns of MCI and NC, except for the M1 module (Fig. 2A). The preservation Z_summary_ score of M1 was 12 in NC, while lower than 2 in AD and MCI, indicating a breakdown of co-expression relationship of M1 neural function in AD and MCI patients (Fig. 3B). The eigengene value and module preservation of AD and MCI exhibited similar patterns in M1, not in other modules (Fig. 2B, 3B). It implies that the overall change of miRNA expression profile mainly occurred after the AD pathology, while neural function related miRNA dysfunction happened in NC-to-MCI stage before the appearance of AD pathological significance.

Additionally, the tissue-specific analysis revealed that 24 out of 26 M1 hub miRNAs were derived from the brain tissue (Table 3). This implies that the exosome secreted by the brain’s disorder had passed through the BBB, resulting in the exosomal miRNA co-expression that could be detected in the peripheral blood. The expression levels of 8 miRNAs decreased simultaneously in both AD and MCI patients (Table 2). All these miRNAs were in M1 module, and six of them, including miR-124-3p, miR-9-5p, miR-9-3p, miR-153-3p, miR-338-3p and miR-125b-5p, were M1 hub miRNAs derived from brain tissue (Table 3, red). Through enrichment analysis, these miRNAs may play post-transcriptional regulator roles in axonogenesis and neural development (Fig. 2D). In previous studies, miR-125b is down-regulated in plasma, serum, and CSF of AD patients^40, 41^. This downregulation is correlated with canonical diagnosis standards, including MMSE scores, Aβ42, T-tau, and P-tau^42^. However, miR-125b hasn’t been shown to change in other neurodegenerative disorders^43^, which indicates the plasma exosome level of miR-125b has the potential to be a non-invasive biomarker for distinguishing AD from other neurodegenerative diseases. Human studies of AD patients also showed a similar decreased expression of miR-9-5p^47^ and miR-9-3p^44^ in CSF and miR-338-3p in plasma exosome. The evidence from these studies suggests that the circulating exo-miRNA levels could reveal the miRNA expression and biological function in the brain.

In 26 hub miRNAs of M1, 15 miRNAs have reported as AD biomarkers or miRNAs with AD-related neural function (Table 3). As a correlation network, the remaining 11 miRNAs in M1 could also play a role in disease progression. In these miRNAs, only miR-770-5p slightly increased in AD, the rest were down-regulated in AD (Table 3). miR-770-5p and miR-708-5p relate to diverse cancers^45^; miR-129-2-3p in plasma is predicted as a non-invasive biomarker for human refractory epilepsy^46^, and miR-218-5p is involved in the pathogenesis of chronic obstructive pulmonary disease^47^. Overall, novel biomarkers from non-invasive blood may represent better diagnostic power for AD, which might lead to a better prognosis.

Ion channel blockade is related to Alzheimer’s amyloid peptide neurotoxicity^48^. In the module preservation analysis, M11 ion channel exhibited model conservativeness above threshold in AD, but not in MCI and NC (Fig. 2). Construction of M11 co-expression revealed the post-translational regulation of ion channel had started in terms of AD onset. The relationship of M11 miRNA co-expression and several health indicators especially HDL suggested that the function change of ion channel might contribute to the pathological of AD patients.

In summary, we compared the miRNAs derived from exosomes between AD, MCI, and NC participants. Using WGCNA method, we constructed 13 coexpression network modules, in which M1 neural function represented a high relationship with AD diagnosis. Module preservation analysis revealed the M1 module was conserved in NC but not in AD or MCI, indicating the dysfunction of post-transcriptional regulation in the neural system. Meanwhile, 8 miRNAs that simultaneously down-regulated in AD and MCI were all in M1 module, and could be biomarkers for AD. The tissue-specific approach further confirmed that the origin of M1 hub miRNA was from the brain tissue. Of 26 hub miRNAs in M1, 15 had been studied closely related to disease pathology, and the other 11 miRNAs were consequently suspected of playing important roles in AD.

## Acknowledgements

We are grateful to those who agreed to donate their blood for research and who participated in the described observational studies. We thank Dr. Yan Zeng for her help of initiating this study. This research was supported by the National Key Research and Development Program of China (No. 2020YFC2002902), Science, Technology and Innovation Commission of Shenzhen Municipality under Grant No. JCYJ20170412153100794 and No. JCYJ20180507183615145, National Natural Science Foundation of China (No. 81941016).

## Author contributions

Y.S., S.R., and designed experiments. Y.S., Z.H., S.Z. and L.Z. carried out experiments. Y.S. and Z.H. analyzed data. W.L., J.D. and T.L. provided advice on the interpretation of data. Y.S., Z.H., W.L., B.L. and S.R. wrote the manuscript with input from co-authors. B.L., Y.D. and S.R. provided tissue samples. W.Z., N.C. and S.R. supervised the study. All authors approved the final manuscript.

---

**Figure 4.** Correlation of miRNA profiles in 13 modules and the clinical traits of AD patients. Upper, Module eigengenes (the same 13 modules in Fig. 2A) were correlated with clinical traits of AD patients. Strength of positive (red) or negative (blue) correlation is shown by two-color heatmap, with *p* < 0.05 values provided. Lower, the measurement of plasma parameters (SIRT1, IL-6, T-TAU, P-TAU, Aβ42) from 26 AD patients.

## Supplementary files

**Supplementary Table 1.** Clinical data of patients with AD and MCI and NC.

**Supplementary Table 2.** Clinical and demographic data used for WGCNA analysis.

**Supplementary Table 3.** Differentially expressed miRNA between AD, MCI and NC.

**Supplementary Table 4.** GO functions of 13 modules.

**Supplementary Table 5.** kME value of each miRNA in 13 modules.

